# Generating Realistic Morphologies of Neurons in Rodent Hippocampus with DCGAN

**DOI:** 10.1101/363481

**Authors:** Darian H. Hadjiabadi

## Abstract

Dendritic size and branching patterns are important features of neural form and function. However, current computational models of neuronal networks use simplistic cylindrical geometries to mimic dendritic arborizations. One reason for this is that current methods to generate dendritic trees have rigid a priori constraints. To address this, a deep convolutional generative adversarial network (DCGAN) trained on images of rodent hippocampal granule and pyramidal dendritic trees. Image features learned by the network were used to generate realistic dendritic morphologies. Results show that DCGANs achieved greater stability^∗^ and high generalization, as quantified by kernel maximum mean discrepancy, when exposed to instance noise and/or label smoothing during training. Trained models successfully generated realistic morphologies for both neuron types, with high false positive rate reported by expert reviewers. Collectively, DCGANs offer a unique opportunity to advance the geometry of neural modeling, and, therefore, to propel our understanding of neuronal function.

∗ A “stable/stabilized DCGAN”, as mentioned throughout this work, is a DCGAN which was stable throughout training.

## 1 Introduction

Neural dendrites are the principal sites for synaptic input; their shapes and sizes across cell classes are highly variable [15]. Furthermore, complexity of dendritic arbors increases during development [17], suggesting that morphology plays a fundamental role in determining neuronal function.

To date, biophysically detailed computational models of the rodent hippocampus has emerged as a tractable tool for investigating open questions in neuroscience. Such models have allowed researchers to gain a deeper understanding of the interneuronal mechanisms underlying theta oscillations [4] and the generation of hyperexcitable networks in the epileptic brain [11]. The predictive power of these tools have [10] and will continue to form a symbiosis with experimental design.

However, one overwhelming problem of current models is that dendritic trees are often modeled using generic cylindrical geometries [11]. As a result, fundamental passive (membrane capacitance, input resistance, electrotonic length) and active (action potential generation and propagation) properties are disconnected from reality. Altogether, this will manifest in inaccurate estimation of electrical attenuation, synaptic integration, and plasticity [7] - processes integral to neuronal function and overarching network dynamics.

Generative methods have emerged as an intriguing opportunity for creating and deploying synthetic dendritic morphologies in computational models. While labs have leveraged these techniques [19], it has not been readily adopted by the field. Particularly, many current algorithms suffer from a priori constraints [3], overall limiting generational diversity. Fortunately, advances in deep learning offer a unique opportunity to generate realistic morphologies by training a model to *learn* useful features.

Here, a deep convolutional generative adversarial network (DCGAN) [14] was trained on thousands of images of granule and pyramidal cell dendritic reconstructions (input) from rodent hippocampus. Features learned by the network were used to output realistic images containing dendritic morphologies. Collectively, DCGANs are in a prime position for advancing state of the art computational models of neuronal networks.

## 2 Related Work

### 2.1 Semi-automated methods

Building geometrically realistic neurons for computational models can be done by tracing. While semi-automatic reconstruction software such as Neuromantic [12] exist, the process is tedious and time consuming [2]. Therefore, considerable efforts have been aimed at building synethetic neurons that statistical similarity to real neurons.

### 2.2 Algorithms for generating neural morphologies

The foundation of modern generative algorithms revolves around the work of Hillman, who described seven fundamental features of neuronal morphology [6]. With this, Asocli and Kirchmar developed the *L*-*Neuron* algorithm that samples from these feature density distributions [3]. Unfortunately, a finite mixture model to the data requires a priori knowledge of the number and types of distributions. Collectively, only a small subset of the total parameter space may be sampled and used for morphological development. Evolutionary algorithms have since emerged as a way to explore broader ranges of this hard coded parameter space [22]. Nonetheless, these parametric methods do not exclude the possibility that other features, not described by Hillman, can be useful towards explaining morphology.

Additional generational methods exist that make use of non-parametric kernel density estimation (KDE) [23], and degree-constrained minimum spanning tree (DCMST) [1]. Data driven non-parametric methods have been used only for few cell classes but have shown promising results [23]. DCMST is an NP-hard problem that operates under the biophysically plausible assumption that neural wiring is minimized for optimizing information transfer. However, it is unclear if this theory holds true for all neurons, and may overall limit generational diversity.

### 2.3 Generational methods in deep learning

Generative models have permeated the field of deep learning. Particularly, variational autoencoders (VAE) [16, 9] and generative adversarial networks (GAN) [5] have showed significant promise. VAEs use statistical inference and are known to train much easier than GANs. However, GAN variants, such deep convolutional GAN (DCGAN) [14], are known to generate higher quality images compared to VAEs. One reason for this is that convolutional neural networks have made remarkable advances in image classification [20]. This is primarily the result of a hierarchical feature extraction architecture that generalizes well to images. Additionally, learned features can be visualized, allowing one to find the approximate purpose of each filter. Further advances towards stabilization, such as training with instance noise [21] and label smoothing [18], make DCGANs a premier tool to generate realistic dendritic trees.

### 2.4 Using deep learning to generate dendritic trees

It is argued that a generative deep learning approach offers a unique advantage to current methods, particularly those with a priori assumptions [3]. Specifically, the parameters of a neural network are mutable. This means that the feature space is learned rather than hard coded, allowing a network to extract a variety of features that may be non-intuitive to humans. Such features may be useful for modeling different classes of neurons, where particular ‘fundamental features’ [6] may or may not have significant influence. To date, no deep network trained to generate dendritic arborizations has been reported.

## 3 Methods

### 3.1 Deep convolutional generative adversarial network (DCGAN)

Generative adversarial networks (GANs) [5] are a class of methods for learning generative models. The GAN can be broken into two distinct neural networks: generator and discriminator. Let *X* = ℝ^*nxn*^ be the space of natural images. The generator can be defined as *G*: *Z* → *X* where *Z* is a latent space. The discriminator will be defined as *D*: *X* → [0, 1). The generator will take as input random noise from latent space *Z* and output an image. The discriminator will then need to decide if the generated image is real or fake. It does so by outputting a probability that correspond to likelihood of an image being drawn from the distribution of real images. The generator’s objective will be to trick the discriminator into thinking a generated image is real. Ultimately, this turns into a minimax game which can be described formally as:

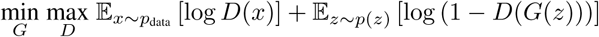

where *z* ~ *p*(*z*) are the random noise samples, *G*(*z*) are the generated images using the neural network generator *G*, and *D* is the output of the discriminator.

The convolutional neural network architectures for the generator and discriminator are shown in **Tables 1 and 2**, respectively. FC = fully connected. Maxpool layers were implemented with 2×2 kernel size and stride of 2. Leaky ReLU units had a negative slope of 0.01. For conv layers in both tables, S=stride; P=padding. In total, the generator network had 4351217 parameters and discriminiator network had 1250241 parameters.

**Table 1:** Generator architecture

| Layer | Volume Dimensions | Num Parameters |
| --- | --- | --- |
| Input | 96x1 | 0 |
| FC | 96x1024 | 99328 |
| ReLU | - | 0 |
| Batchnorm | - | 2048 |
| FC | 1024x4096 | 4198400 |
| ReLU | - | 0 |
| Batchnorm | - | 8192 |
| ConvTranspose | 64x32x4x4 (S2;P1) | 32800 |
| ReLU | - | 0 |
| Batchnorm | - | 64 |
| ConvTranspose | 32x16x4x4 (S2;P1) | 8208 |
| ReLU | - | 0 |
| Batchnorm | - | 32 |
| ConvTranspose | 16x8x4x4 (S2;P1) | 2056 |
| ReLU | - | 0 |
| Batchnorm | - | 16 |
| ConvTranspose | 8x1x3x3 (S1;P1) | 73 |
| Tanh | 1x1 | 0 |

**Table 2:**
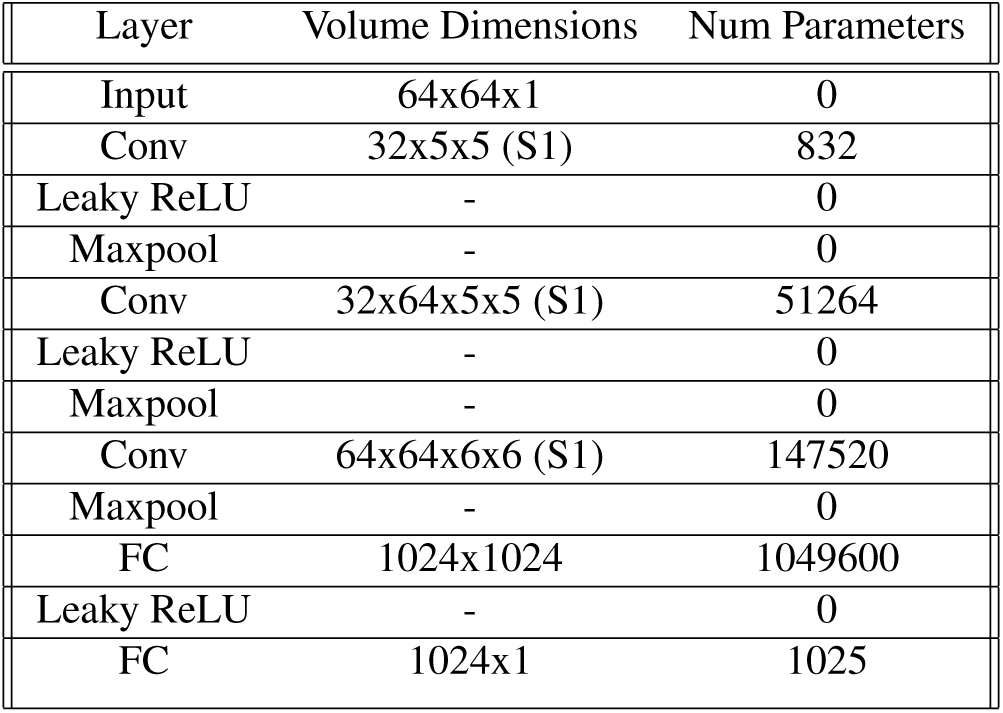
Discriminator architecture

### 3.2 Loss function and optimization

Training GANs requires finding a Nash equilibrium of a non-convex game, a non-trivial problem. Instead, gradient descent techniques are used to minimize a loss function. The discriminator will seek to maximize the probability that it can make a correct choice on both real and fake data. The discriminator loss can therefore be formalized as:

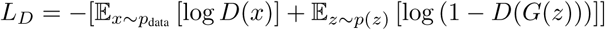

The generator will seek to maximize the probability of the discriminator making the incorrect choice on generated data. The generator loss can therefore be formalized as:

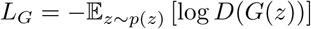

In this work, ADAM [8] was used to minimize the loss functions.

### 3.3 Stabilization

GAN training is notoriously unstable. To address these concerns, numerous ‘tricks’ have been developed. In this work, label smoothing [18] and instance noise [21] were leveraged. A nice graphical representation of the two methods can be found here: http://www.tinyurl.com/znnnvej.

To motivate the nature of these methods, it is worth diving deeper into the internals of a GAN. The loss function of the discriminator is the negative expectation of the log likelihood ratio 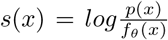 where *f_θ_* represents the generative model and *p* is the real data. One major assumption underlying optimization convergence is that *s*(⋅) is finite. For this condition to hold, the support (i.e. inputs not mapped to 0) must overlap. *f_θ_* is degenerate by construction, and the training process attempts to build a model that overlaps with *p*. The stabilization techniques below attempt to foster the process of generating support overlap by altering the log likelihood function. In a sense, the discriminator is ‘tricked’.

#### 3.3.1 Label smoothing

In this technique, labels of some real data sample *x* ~ *p* are flipped so that the discriminator is trained to think they are from *f_θ_*. If this is performed with probability *ν*, the log likelihood function is now 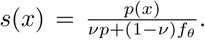 Since the denominator now includes information regarding the distribution from both the real and generated data, support overlap is more likely to occur. The structure of the logistic loss landscape is not altered dramatically, however, and so non-convergence may still occur.

#### 3.3.2 Instance noise

Here, noise is input into both the generated and fake data. White Gaussian noise *ϵ_σ_* is convolved with images so that the log likelihood function is now 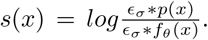 Because the noise distributions overlap, *s*(⋅) will ‘behave’ better.

### 3.4 Kernel maximum mean discrepancy (MMD)

Evaluation metrics for GANs have been thoroughly researched. Kernel maximum mean discrepancy (MMD) has been recently shown to assess overfitting [24]. Kernel MMD can be thought of as a measure between the statistical dissimilarity between two datasets (i.e. higher MMD means datasets are not similar) and can be written as:

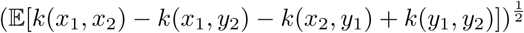

where *x*_1_, *x*_2_ are drawn from dataset *X*, *y*_1_, *y*_2_ are drawn from dataset *Y*, and *k* is a Gaussian kernel with width *σ*;, written as:

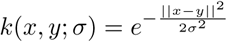

In practice, the kernel MMD value can be approximated as:

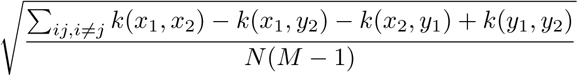

where *N* = ∣*X*∣ and *M* = ∣*Y*∣. To assess overfitting, the MMD metric is calculated between training data *X* and generated data *F* - call this *ρ*(*X*, *F*). Next, the MMD metric is calculated between validation data *Y* and generated data *F* - call this *ρ*(*Y*, *F*). The ‘metric gap’ is defined as:

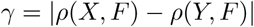

Higher values of *γ* imply overfitting. Furthermore, kernel MMD is used in this work to compare between models. The rational here is that if two models display low MMD gap (i.e. don’t overfit) but model A has a lower post-training *ρ*(*X*, *F*) compared to model B, then model A is superior.

### 3.5 Qualitative assessment

To assess if generated images ‘looked’ real, 64 fake and 64 real images were pseudorandomely shuffled and blurred with 5 x 5 mean smoothing. The 128 image stacks were given to 5 experts (PhD holders) in the field of neuroscience who rated each image as 1 if real or 0 if fake. This was done for both granule pyramidal cells separately.

## 4 Dataset

All granule and pyramidal cell reconstructions from rodent hippocampus were extracted from http://www.NeuroMorpho.org. NeuroMorpho is an well recognized and standardized database containing neural reconstructions submitted by labs around the world. 3107 granule and 5708 pyramidal cell reconstructions were downloaded for processing. Point cloud files were ingested, and dendritic points extracted and centered. Any point cloud files lacking dendritic trees or with sparse reconstructions were excluded. Sparse reconstruction was determined if a dendritic cloud had less than 100 points, an admittedly arbitrary threshold. Lastly, aligned clouds that passed both conditions were resampled to output images of size 64 x 64 pixels. These 64 x 64 pixel images were fed into the DCGAN.

Ultimately, 2802 granule and 5287 pyramidal cells passed both conditions. 90% of images from each class were used for training (2522 granule, 4759 pyramidal) and the remaining 10% for validation (280 granule, 528 pyramidal). **Figure 1** shows example reconstructions and 64 × 64 pixel image outputs for both neuron classes. It can be observed that morphologies are unique between the two classes.

**Figure 1:**
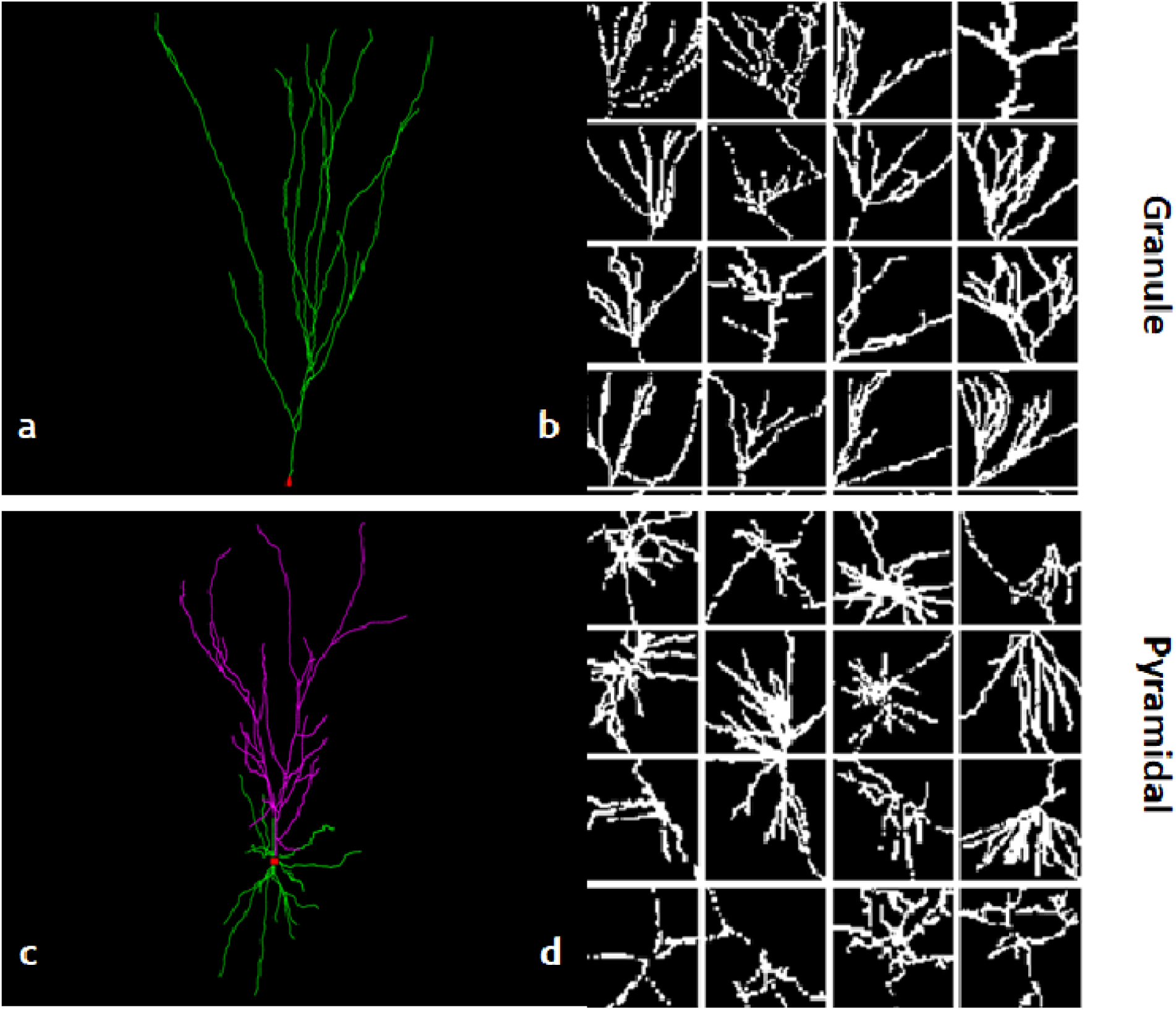
(a, c) Example neuron reconstructions of granule (a) and pyramidal cell (c). (b,d) Example granule (b) and pyramidal (d) 64 x 64 arborization images output from preprocessing.

## 5 Experiments

### 5.1 Hyperparameters

Separate DCGAN models were trained for granule and pyramidal cells with PyTorch [13]. The Adam optimizer [8] was used for all training with fixed hyper-parameters (*β*1, *β*2) = (0.5, 0.999) and *ϵ* = 10^−8^ for numerical stability. Adam was used because it incorporates advantages from momentum based and adaptive optimization techniques. Training was limited to mini-batch size of 64 due to available computational resources. For label smoothing, probability of label flipping was set to 0.1. This was implemented by swapping the ‘fake’ and ‘real’ logits. For instance noise, Gaussian white noise with *μ* = 0 and initial *σ* = 10^−2^ was input to images. *σ* underwent linear decay, with updates after each epoch, to 0 during the training process.

To find the optimal learning rate hyper-parameter, 20 values randomly picked from the range [10^−5^, 10^−2^] were tested. Qualitative assessment of image quality and loss curves suggested that 5.94∗10^−4^ and 6.85∗10^−4^ are ideal candidates for granule and pyramidal DCGAN, respectively (**Figure 2**).

**Figure 2:**
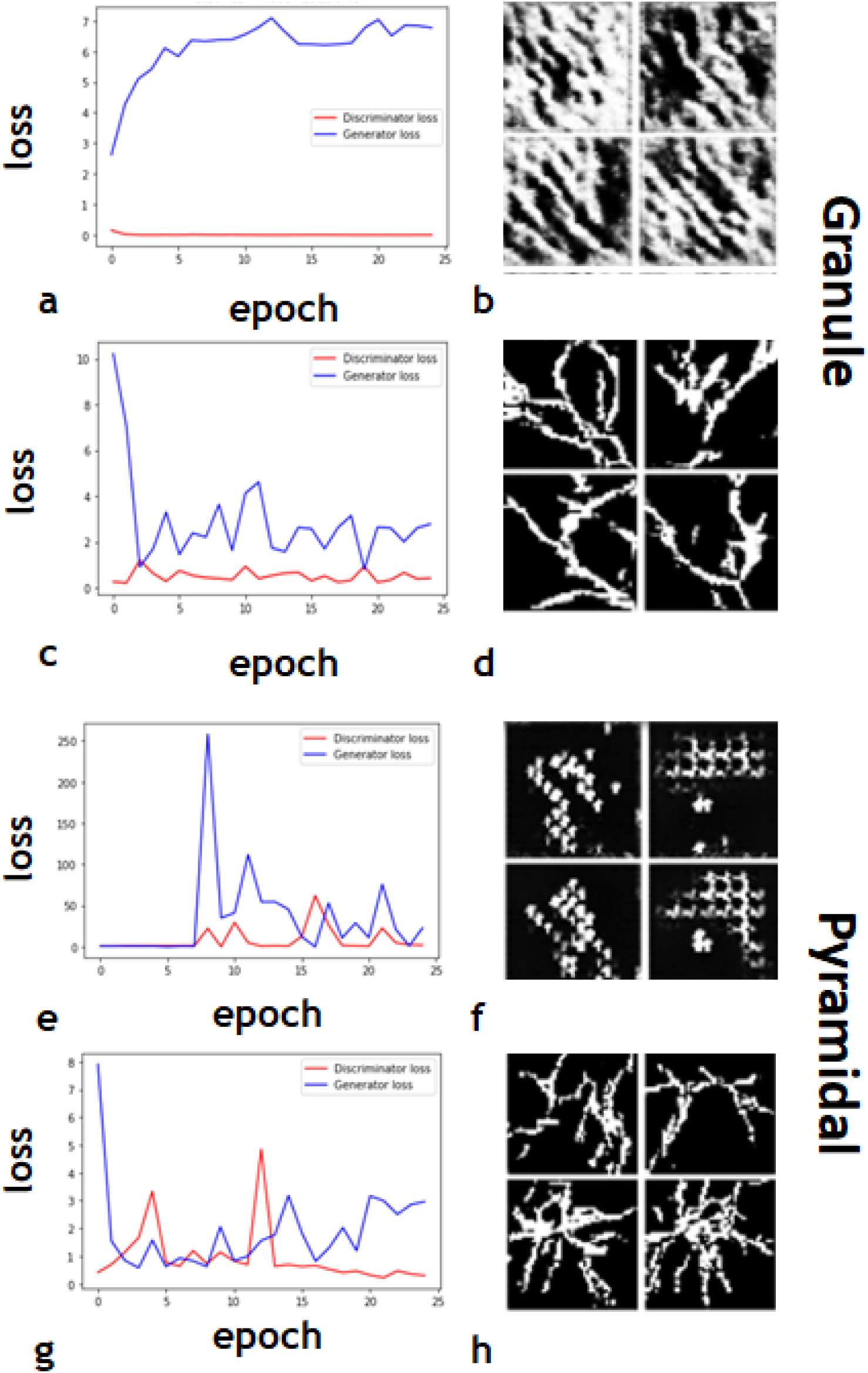
Learning rate hyperparameter search. (a,b) An example of poor training on images of granule cells (learning rate 2.26 ∗ 10^−5^). (c,d) Successful training of DCGAN on images of graunle cells (learning rate 5.95 ∗ 10^−4^). (e,f) same as (a,b) but for pyramidal cells and learning rate 4.33 ∗ 10^−3^. (g,h) same as (c,d) but for pyramidal cells and learning rate 6.85 ∗ 10^−4^.

### 5.2 Label smoothing and instance noise stabilized DCGANs during training

Models were then trained with the above mentioned learning rates for 100 epochs. DCGAN trained without stabilization techniques exhibited generator loss that slowly increased over the course of training (**Figure 3**). This furthermore resulted in poor quality generated images.

**Figure 3:**
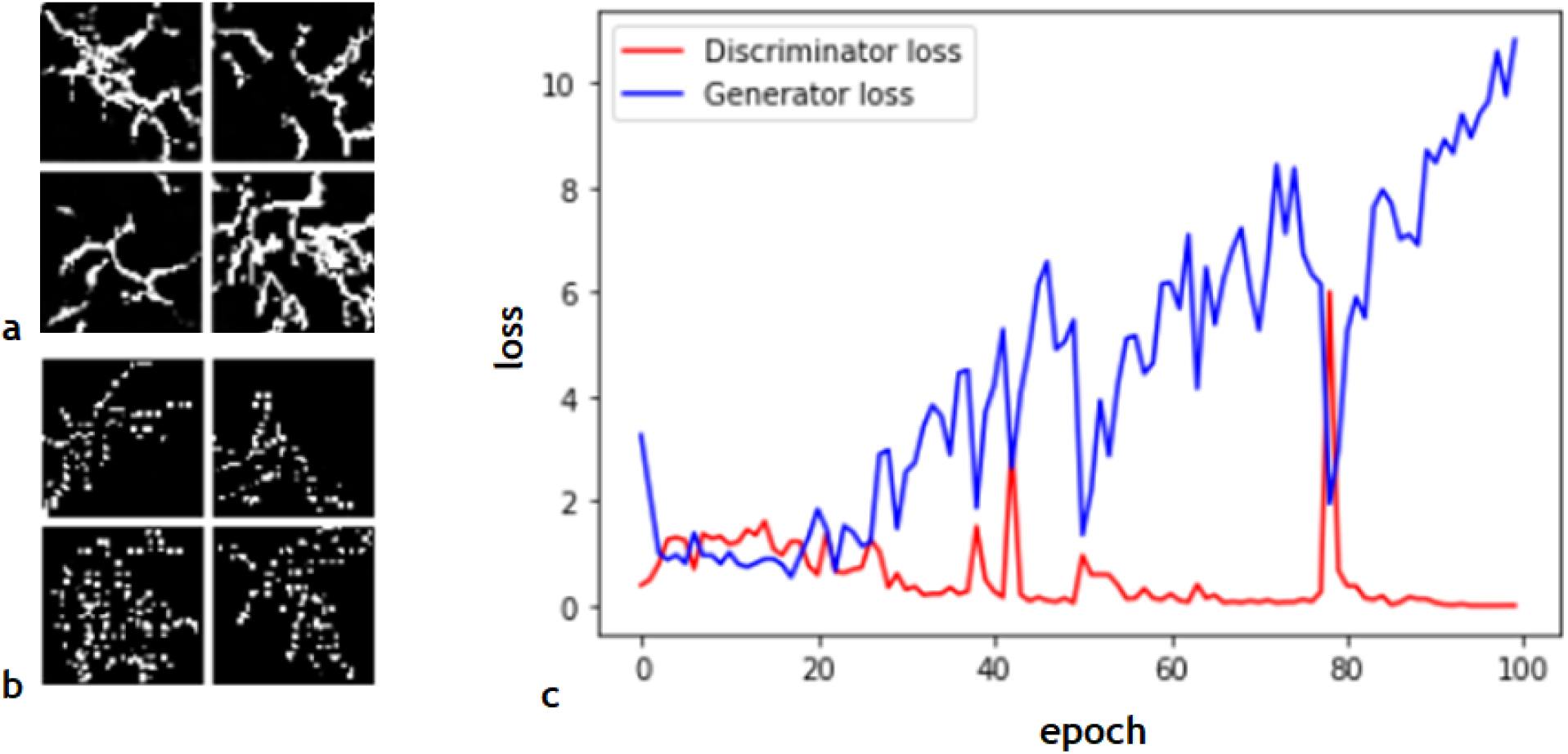
DCGANs can be unstable over long training durations. (a) Output of DCGAN trained for 25 epochs on images of pyramidal cells and learning rate 6.85 ∗ 10^−4^. (b) Output of DCGAN after 100 epochs of training on the same learning rate. (c) Discriminator (red) and generator (blue) loss curve over training. Notice the slow increase in generator loss over training.

To stabilize the models, a combination of instance noise and label smoothing was applied (see methods). Kernel mean maximum discrepancy (MMD) (see methods) between 256 generated and 256 training images was used to compare models after 100 epochs of training. Low kernel MMD would indicate high statistical similarity between the two datasets, an ideal situation. For granule cells, models trained for 100 epochs with both instance noise and label smoothing resulted in the lowest kernel MMD over multiple Gaussian kernel widths (**Figure 4a**). For pyramidal cells, this was true whether instance noise and/or label smoothing was used (**Figure 4c**). For both cell types, generator loss stabilized and discriminator loss exhibited periodic spikes (**Figures 4b,d**).

**Figure 4:**
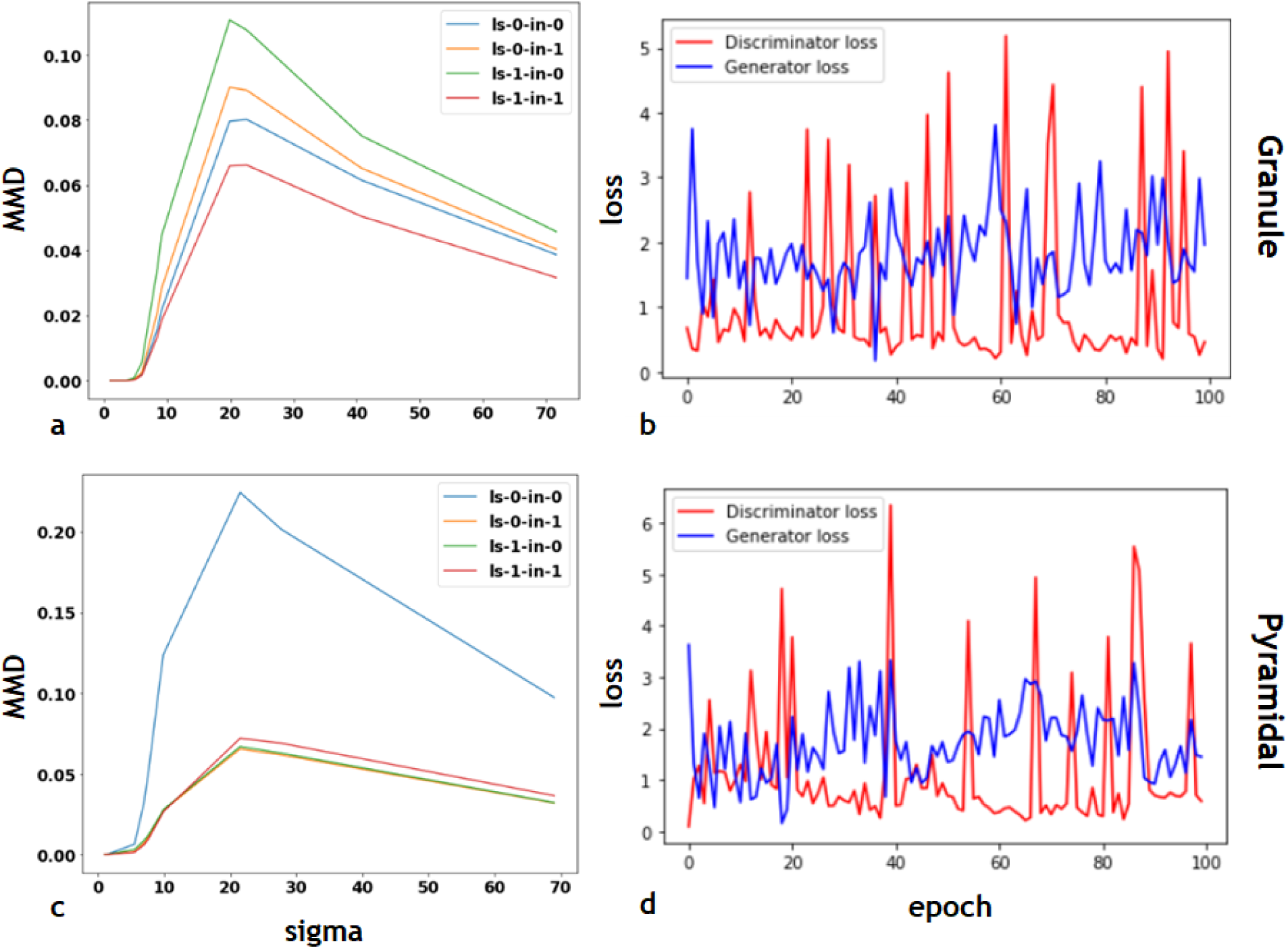
Label smoothing and instance noise help stabilize DCGAN training. (a) Plots of kernel MMD between training data and DCGAN generated images over varying Gaussian kernel widths. DCGAN model was trained on granule cells with learning rate 5.94 ∗ 10^−4^. One can see that the red curve (label soothing on, instance noise on) has lowest kernel MMD. (b) Plot of discriminator (red) and generator (blue) loss over training duration for DCGAN model trained with label smoothing and/or instance noise. (c) Same as (a) except for DCGAN trained on pyramidal cells with learning rate 6.85 ∗ 10^−4^. (d) Discriminator (blue) and generator (red) loss curves for DCGAN trained with label smoothing. Legend key: ls = label smoothing. in = instance noise. 0 = off. 1 = on.

### 5.3 Stabilized DCGANs were well-generalized

Low kernel MMD can imply overfitting if the synthetic images simply memorized the training images. To test for this possibility, kernel MMD with *σ* = 25.0 was calculated between (1) 256 training images and 256 generated images; (2) 256 validation images and 256 generated images. A large difference between (1) and (2) - i.e. the metric gap (see methods) - is an indicator of overfitting. The above mentioned sigma value was chosen because kernel MMD reported highest values as seen in **Figures 4a,c. Figures 5a,c** shows that low gap values were calculated for both granule (< 0.01) and pyramidal cells (< 0.02) across all training epochs. However, pyramidal cell training appeared to be slightly more unstable. Furthermore, it was hypothesized that kernel MMD would start high and decrease over training as the generative model learned to output realistic images. **Figures 5b,d** confirms this, ultimately providing quantifiable evidence that training improved generation quality.

**Figure 5:**
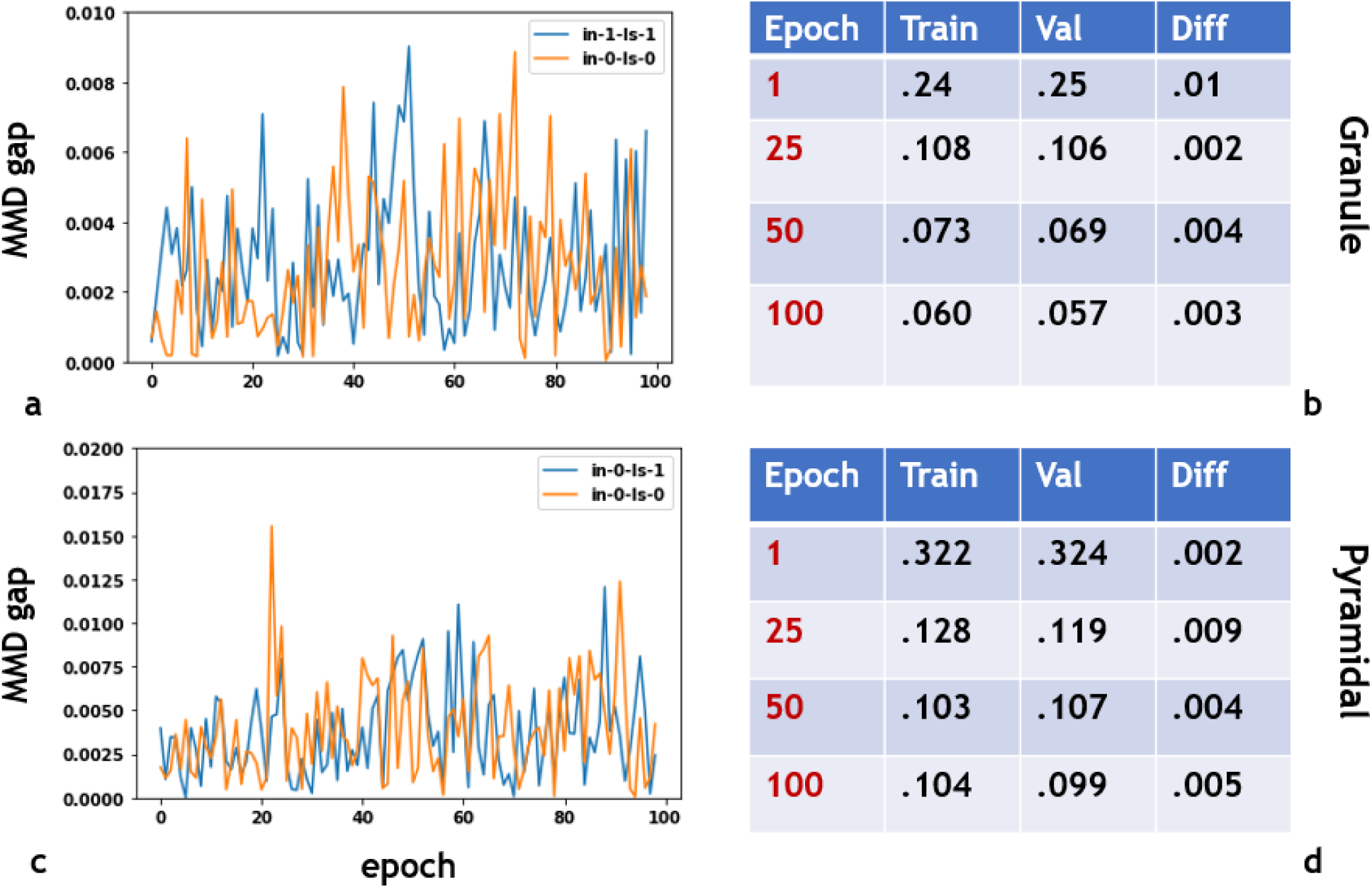
DCGAN models with label smoothing and/or instance noise do not overfit training data. (a) Plot of kernel MMD metric ‘gap’ (see method) for DCGAN model trained on images of granule cells. Blue curve represents model trained with label smoothing and instance noise. Orange curve represents vanilla DCGAN. (b) Table of gap values for DCGAN model trained on granule cells with label smoothing and instance noise. (c) Pyramidal cell DCGAN equivalent of (a). Blue curve represents model trained with label smoothing on. Orange curves represents vanilla DCGAN. (d) Table of gap values for DCGAN model trained on pyramidal cells with label smoothing.

### 5.4 Visualizing convolutional layers

Filters from the first convolutional layer of the discriminator and generator were visualized (**Figures 6a-d**). It can be seen that generator filters attempted to learn edge orientations and possible bifurcations (**Figures 6a,c**). Furthermore, it is evident the that discriminator filters attempted to dissect complicated branching structures (**Figures 6b,d**). To further investigate how label smoothing and/or instance noise affects network structure, filter distributions were plotted between models trained with or without stabilization (**Figures 6e,f**). Distributions appeared almost identical for most layers with the exception of third convolutional layer of the generator and first convolutional layer of the discriminator.

**Figure 6:**
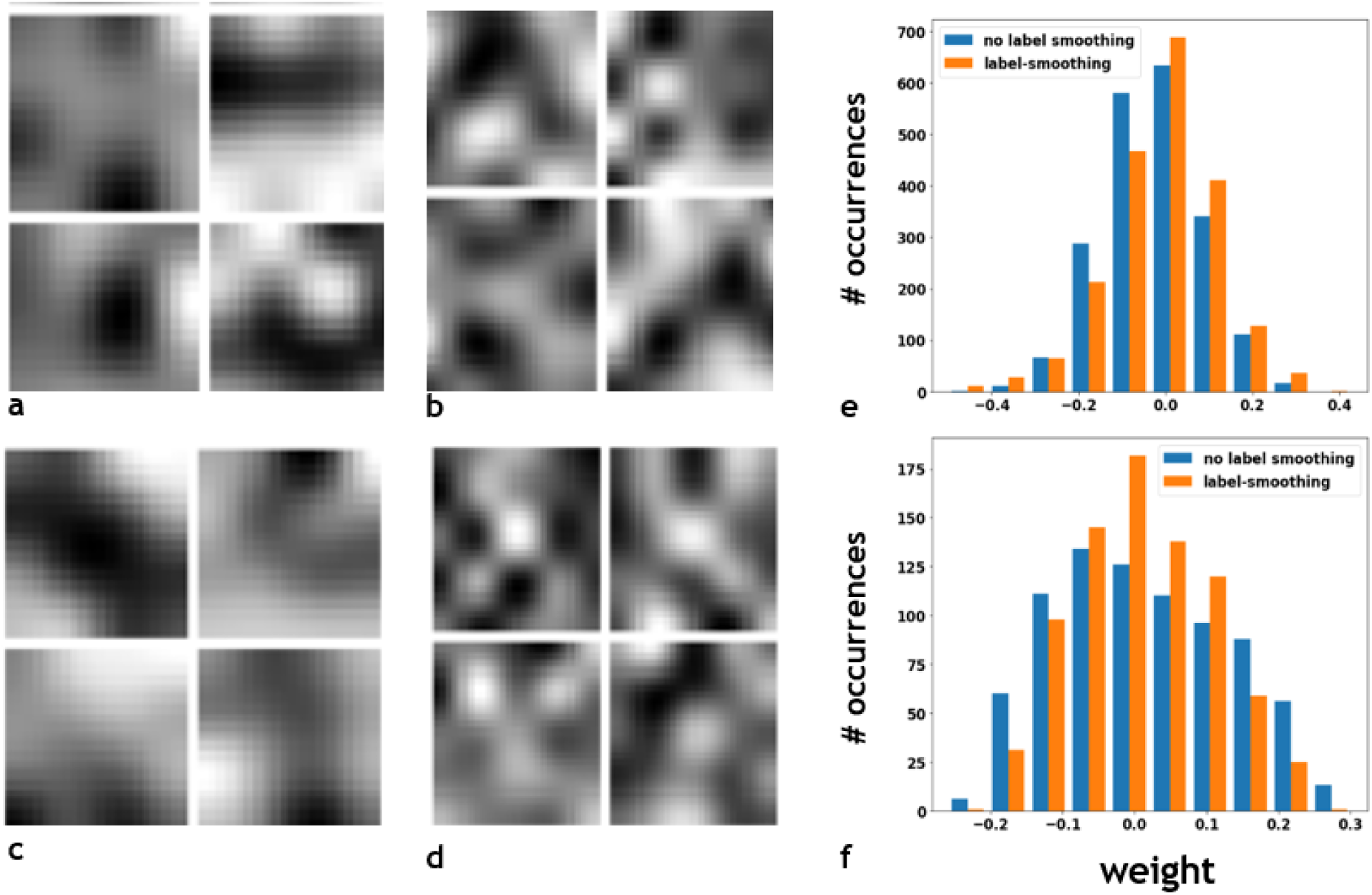
(a,c) Examples of filters from the first convolutional layer of DCGAN generator trained on images of granule (a) and pyramidal (c) cells for 100 epochs. (b,d) Examples of filters from first convolutional layer of DCGAN discriminator trained on images of granule (b) and pyramidal (d) cells for 100 epochs. (e) Comparison of weight distribution in third convolutional layer of generator for DCGAN model trained with (orange) and without (blue) label smoothing. (f) Similar to (e) but for the first convolutional layer of discriminator.

### 5.5 Trained DCGANs generated high quality dendritic morphologies

The best granule and pyramidal DCGAN models (see 5.3) were used for dendritic morphology generation (**Figure 7**) and qualitative evaluation. Shuffled datasets containing real and fake morphologies (**Figure 8**) were presented to expert reviewers. Results (**Table 3**) show false positive rates of 0.54 and 0.44 for granule and pyramidal cells, respectively. That is, almost half of the generated images were considered real to the expert eye. However, true positive rate was 0.52 for granule and 0.48 for pyramidal cells, suggesting difficulty in identifying features of true morphologies. Overall accuracy was approximately 50% for both classes of cells.

**Figure 7:**
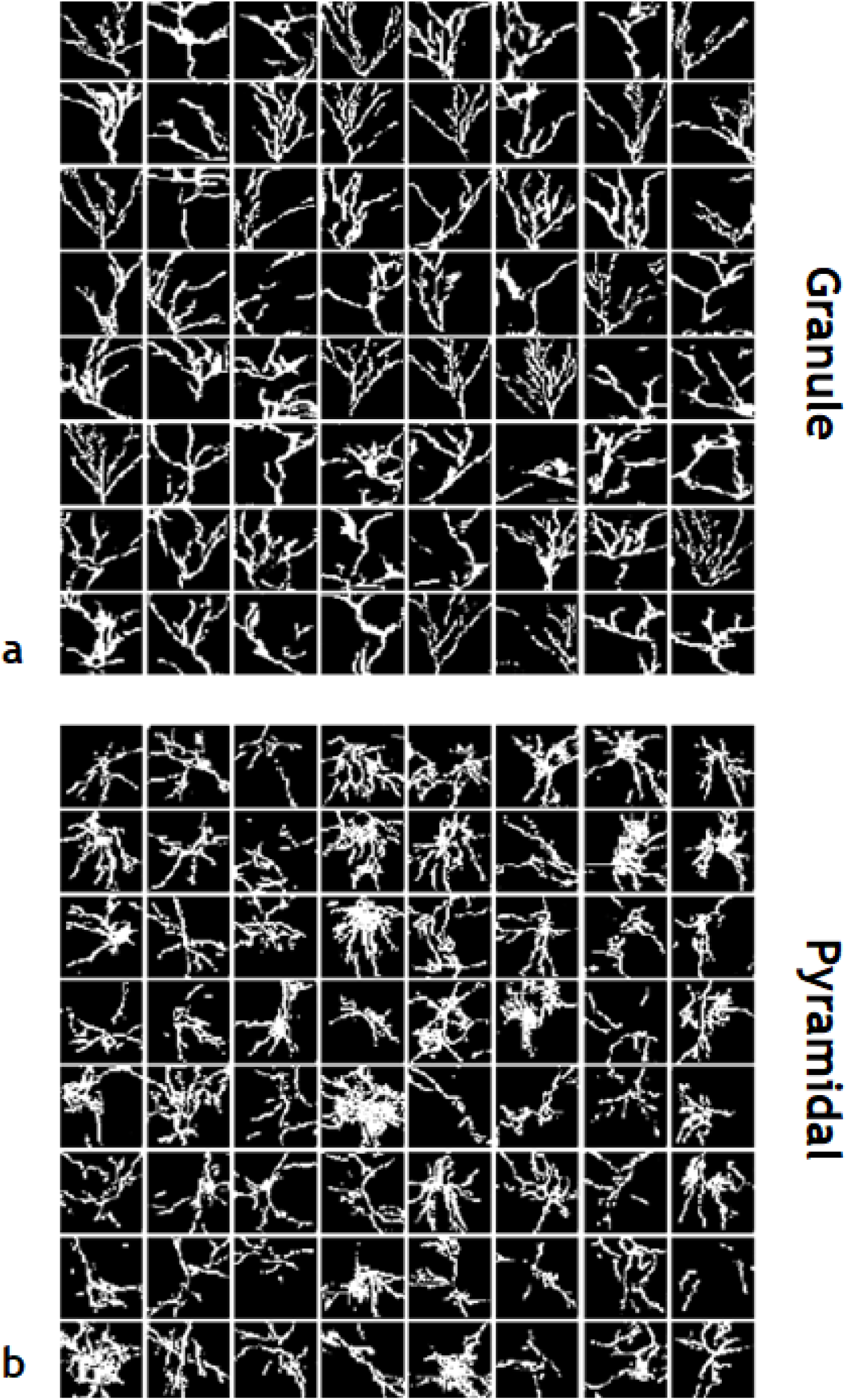
Generated images of granule (a) and pyramidal (b) cell dendritic arborizations from best trained DCGAN models.

**Figure 8:**
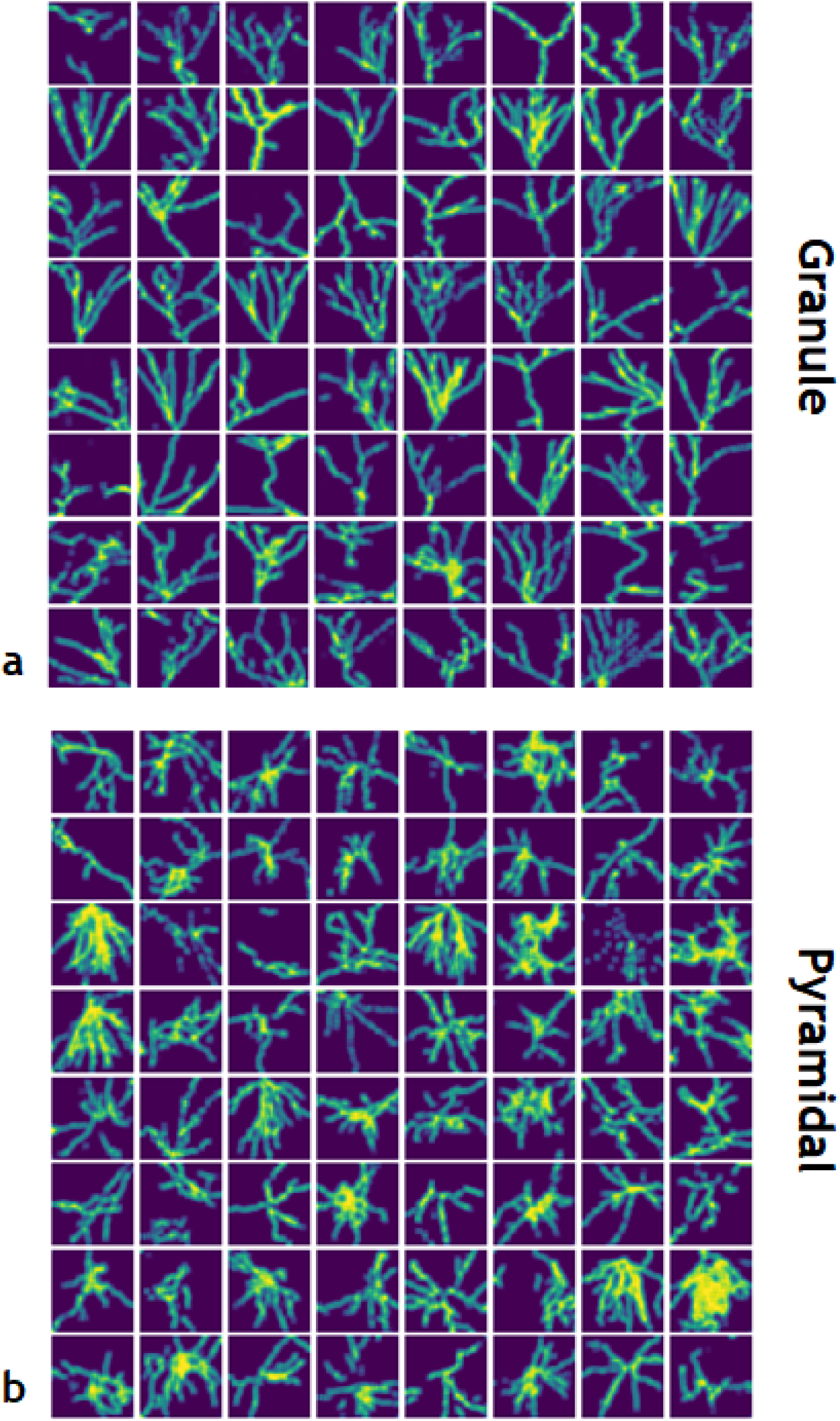
Expert reviewers were tasked with discerning real vs fake. (a) Generated images of granule cell dendritic arborizations pseudorandomely shuffled with training data. (b) Generated images of pyramidal cell dendritic arborizations psuedorandomely shuffled with training data. Images were blurred with 5 x 5 spatial mean filter. Can you discern real from fake?

**Table 3:** Results of expert assessment to discern real vs fake granule and pyramidal cell dendritic morphologies. n = 5 for both cell classes. True positive represents a correct identification of real image. False positive represents identifying generated image as real. Values represented as mean with standard deviation in parenthesis.

| Statistic | Granule | Pyramidal |
| --- | --- | --- |
| True positive | 0.52 (.18) | 0.48 (0.15) |
| False positive | 0.54 (0.19) | 0.44 (0.11) |
| Accuracy | 0.49 (0.14) | 0.53 (0.12) |

## 6 Discussion

### 6.1 Stabilization techniques and its affect on network infrastructure

It is evident that application of label smoothing and instance noise resulted in stable and well-generalized DCGAN models. High frequency spikes in discriminator loss are expected given the nature of these techniques are to ‘trick’ the discriminator. However, inspection of weight distributions across discriminator and generator convolution layers was insufficient to isolate the affect of label smoothing and/or instance noise on network infrastructure. No identifiable trends were observed for the granule DCGAN despite observation of low frequency increase in generator loss over training. Furthermore, the weight distributions presented for pyramidal DCGAN is not convincing evidence towards any particular conclusion.

### 6.2 Advances over current generative methods

The parameters of a DCGAN model are free to learn whatever features are necessary for the generator to output realistic morphologies. This results in higher-order image features that humans may not have the insight to concept. In this regard, the method proposed here is an advance over current techniques that sample from a pre-defined feature space [3]. Overall, this may lead to increased diversity in generated morphologies, but a more quantitative assessment will need to be performed.

### 6.3 Limitations of this study

Despite the positive outcome, this study has its flaws. It was reported here that DCGAN models generalized well, and therefore can produce images not seen in the training set. However, a comparative study quantifying differences in diversity between the method proposed here and current generational techniques [3] needs to be performed. Secondly, during hyper-parameter search, the ‘best’ learning-rate was identified by qualitative inspection of generated morphologies. While certainly some models can be thrown away from eye-ball inspection, kernel MMD should have been used to isolate the ideal hyper-parameter amongst models with similar looking outputs. Lastly, it is clear from the low true positive rate amongst expert reviewers that there might have been ambiguity in what a real dendritic tree looked like. This might have been caused by downsampling dendritic clouds to 64 x 64 pixel images. Nonetheless, this work shows that generating high quality realistic morphologies is possible via a deep learning approach.

## 7 Conclusion

Computational models of neural networks have become increasingly important for investigating topics in neuroscience. Unfortunately, most models do not attempt to incorporate realistic dendritic trees or use techniques that limit diversity through a priori assumptions. This results in simulations that may not properly model neuron function.

To address this, a deep learning approach was applied to generate realistic dendritic morphologies. Specifically, DCGAN models were trained on thousands of granule and pyramidal cell reconstructions from rodent hippocampus. Results show that trained models are highly sensitive to learning-rate and are unstable unless label smoothing and/or instance noise is implemented. For granule cells, it was shown that both instance noise and label smoothing resulted in the lowest statistical discrepancy between generated and training data, as quantified by kernel maximum mean discrepancy (MMD). This was the case for pyramidal cells whether instance noise or label smoothing was used. Additionally, stable DCGAN models exhibited a high degree of generalization. Lastly, generated images from trained models were of high quality as determined by expert reviewers. Collectively, it is shown that well-engineering DCGANs presents a unique opportunity to add geometric realism to current computational models.

Future work will seek to build 3D morphologies. Furthermore, the diversity between traditional generation techniques and this proposed method will be quantitatively compared.

NeuroGAN public repo can be found here: http://www.github.com/dhadjia1/NeuroGAN

## 8 Contributions and Acknowledgments

D.H. acquired all data, performed preprocessing, built models, performed experiments, and wrote the paper. I.S., G.S., J.F., A.M., and I.R. are thanked for assessing quality of generated morphologies. J.F., A.M., and I.R. provided useful discussion.

